# Neural tube organoid generation: a robust and reproducible protocol from single mouse embryonic stem cells

**DOI:** 10.1101/2024.06.20.599764

**Authors:** Teresa Krammer, Elly M. Tanaka

## Abstract

The development of mammals is a highly complex process, characterized by the necessity for precise concentration- and time-dependent signaling for correct pattern formation and morphogenesis. Despite considerable technological advancements and knowledge gathered, numerous aspects of mammalian development remain elusive. When examining the entire organism, it becomes challenging to disentangle the effects of individual pathways or the mechanism by which external stimuli guide the interference of surrounding tissues and factors. In addressing this complexity, three-dimensional (3D) *in vitro* models such as organoids have emerged as valuable tools. Organoids, derived from embryonic stem cells (ESCs) or induced pluripotent stem cells (iPSCs), exhibit tissue-like features that closely resemble their *in vivo* counterparts in terms of expression patterns and functionality. Importantly, they offer accessibility for manipulation and extensive biological studies within a controlled experimental setting. Despite originating from pluripotent cultures, organoid systems often exhibit heterogeneity and substantial variability, limiting their utility when studying complex and intricate biological questions. Therefore, there is a pressing need for detailed protocols aimed at harmonizing procedures that result in high-quality reproducible data, reduction of materials used and which importantly permit the investigation of convoluted phenomena. In this context, we present an optimized protocol for the cultivation of neural tube organoids (NTOs) *in vitro*. By producing stable culture conditions and offering comprehensive troubleshooting strategies, this protocol enables the reliable and reproducible generation of NTOs which serve as an adequate model to study relevant scientific questions.

**SUMMARY:** Three-dimensional neural tube organoids (NTOs) derived from mouse embryonic stem cells are valuable tools to study the central nervous system during early development. Here, we present a step-by-step demonstration of an optimized protocol for cultivating NTOs *in vitro*, providing stable culture conditions and troubleshooting strategies for reliable and reproducible NTO generation.

## INTRODUCTION

Mammalian development, starting from a single totipotent stem cell, is a complex process that involves the induction of pattern formation and morphogenesis. This development requires tightly regulated cellular programs to manage dynamic interactions between cells and their environments. Stem cells have the intrinsic ability to sense, integrate, and respond to systemic and local signals including morphogen gradients ^1^, mechanical boundaries ^2^, cellular proliferation, and environmental remodeling ^3^. External spatiotemporal cues also drive multicellular responses, initially, in homogenous tissues, to ensure the precise formation of complex structures. Stem cells, distinguished by their self-renewal ^4, 5^ and differentiation capabilities, play a crucial role in embryonic development and adult tissue repair, with main types including embryonic stem cells (ESCs), adult stem cells (ASCs), and induced pluripotent stem cells (iPSCs).

ESCs are extracted from the inner cell mass (ICM) of a blastocyst and are typically cultured in a two-dimensional (2D) environment on feeder cells or extracellular matrix using specific culture media to maintain their pluripotency ^6^. In recent years, the need for more realistic cell culture conditions and technologies increased drastically, thus three-dimensional (3D) cell and tissue models were established to study fundamental biomedical processes. Especially in developmental biology, these 3D models such as organoids offer a robust method for modeling tissue morphogenesis and organogenesis *ex vivo* ^7^. Organoids form through self-organization of proliferating stem cells that spontaneously develop into complex 3D structures through symmetry breaking and pattern formation. This process is similar, though not identical to what is observed *in vivo*, and mostly occurs without external guidance but driven by internal dynamics and interactions ^8^. Since organoids closely mimic the structure and function of actual organs, they can provide a more accurate representation of mammalian biology compared to traditional 2D cell cultures. In additon, one of the primary benefits is the ethical advantage, as organoids reduce the need for animal models, addressing concerns related to animal testing. Furthermore, organoids offer a platform for high-throughput drug screening, improving the prediction of drug efficacy and toxicity. However, there are also several challenges associated with using organoids. While they are more realistic than 2D cultures, they do not fully replicate the complexity and interactions of entire organs within a living organism. Additionally, there can be significant variability observed between organoids, even when derived from the same type of cells, leading to inconsistencies in experimental results. Thus, culturing organoids in a robust and reproducible manner is essential, and requires advanced technical skills, specialized equipment and detailed protocols.

The nervous system is a complex network responsible for coordinating actions and processing sensory information by sending and receiving signals throughout the body, thereby controlling and coordinating all required functions, including movement, sensation, thinking, and autonomic functions such as heartbeat and digestion. The precursor for the central nervous system, consisting of the brain and spinal cord, is the neural tube. It is crucial to understand how neural structures form and function within early neural tube development, to obtain insights into various neurological disorders and developmental abnormalities. Neurodevelopmental disorders are challenging to study *in vivo*, especially when they originate from human subjects due to ethical concerns. Neural organoids such as brain organoids ^9^ have significantly enhanced our understanding of nervous system development and disease by providing versatile, accessible models that closely mimic human brain tissue. These three-dimensional structures, derived from pluripotent stem cells (PSCs), replicate various aspects of brain development, including the formation of distinct brain regions, cellular diversity, and complex neural networks ^10, 11^. By allowing researchers to observe developmental stages in real-time and study key regulatory mechanisms, these organoids offer deep insights into normal brain development. Brain organoids also enable the modeling of neurodevelopmental ^12^ and neurodegenerative diseases ^13^, such as microcephaly ^10, 14^, Zika virus infection ^15^, autism spectrum disorders ^16^ and others, elucidating disease mechanisms and identifying potential therapeutic targets. Additionally, brain organoids serve as valuable platforms for drug testing and development, facilitating the assessment of drug efficacy and toxicity in a human brain-like environment and accelerating personalized medicine approaches. Moreover, they provide a controlled setting to study gene-environment interactions, enhancing our understanding of complex diseases resulting from the interplay of genetic and environmental factors. However, since neural organoids, like brain organoids, are typically formed from aggregates of cells, they often exhibit significant cellular heterogeneity and variability in experimental outcomes, which in turn influences the interpretability of the generated data, posing a considerable challenge for researchers. For this reason, protocol optimization and precise quality control criteria are critical for the attenuation of valuable data that adequately represent the *in vivo* condition.

Meinhardt and colleagues ^17^ made a crucial impact by first reporting a clonal neural organoid protocol mimicking early neural tube development, formed from single embryonic stem cells (ESCs) that drastically minimizes inter-organoid heterogeneity. This innovation allows researchers to study developmental processes and mechanisms, as well as neural patterning, in a precise and defined environment with exceptional resolution. Until now, many laboratories have developed neural organoids, mostly of mouse or human origin, that mimic different parts of the neural tube and its development, as summarized in ^18^. However, to our knowledge, the protocol established by Meinhardt et al. ^17^ is the only method where NTOs reliably form from a single cell, self-organize and efficiently pattern into dorsoventral fates following a single, global pulse of retinoic acid (RA) without the addition of other ventralizing factors such as Sonic Hedgehog Agonist (SAG)^19^, or the usage of microfluidic chambers that provide spatial information due to gradient formation. The protocol provided here was adapted from Meinhardt et al.^17^ as provided in Krammer et al.^19^. In this protocol, single mouse ESCs are seeded in Matrigel and exposed to defined neurodifferentiation conditions. Within two days, these cells clonally expand and epithelialize, forming a 3D round-to-ellipsoid structure with a single lumen and apically-basally stratified cells surrounding it. On day 2, a globally applied 18-hour pulse of the signaling molecule RA enhances neural differentiation, posteriorizes the NTOs from midbrain to hindbrain/cervical spinal cord levels, and induces the formation of a single, functional floorplate by day 6, which patterns the NTO from ventral to dorsal.

The protocol has the following advantages:

- Robust epithelialization within first 3 days,
- Relatively small spherical tissues (∼200 μm) with a single lumen,
- Clonal growth from single cells,
- Serum-free, chemically defined culture medium,
- Gradual neural differentiation from pluripotent cultures within 6 days,
- ‘By-default’ anterior midbrain identity by day 6,
- Dorsoventrally patterned hindbrain/ cervical spinal cord-like structures within 6 days ^19^.

Several aspects have been adapted and differ from the original protocol ^17, 20^ in the following ways:

- Different ESC culture condition,
- Less input cells,
- Improved differentiation medium,
- Changed differentiation protocol (no NOG d0-2) (already used in ^19^),
- Seeding (density) adapted to multi-well plate (96) usage for high throughput analysis leading to 40% increase in NTO formation and patterning efficiency (from initially ∼40% to now ∼80%).

Here, we provide a detailed description of an optimized protocol for a robust and reproducible generation of NTOs in Matrigel for:

- NTO handling in all stages from seeding ESCs for differentiation assay, to maintenance, fixation and analysis.
- Quality control by visual assessment of morphology of NTOs on daily basis.
- Critical characterization of correct NTO identity via antibody staining.

In addition, we offer comprehensive troubleshooting approaches, allowing a critical assessment in every step of the protocol, which is necessary to ensure the reproducibility and reliability of experimental protocols. These detailed instructions help to identify and resolve potential issues, minimize errors, and optimize outcomes. This is crucial for maintaining the integrity of scientific experiments and enabling other scientists to successfully replicate the work and draw comprehensive conclusions from the data generated, thereby advancing the field.

### PROTOCOL

#### General statements

- Cells and NTOs are checked daily for quality assessment before any further procedure (medium replacement, treatment, fixation, etc.).
- The morphology of the NTOs is crucial and can provide significant insights into the viability of the culture and the efficacy of the experiment.
- All reagents (Medium, PBS, Accutase) are pre-warmed at a minimum of 21 °C and maximum 37 °C before use.
- When preparing medium (N2B27, N2B27diff) glutamine is kept for a maximum of 5-10 minutes at 37 °C, better at room temperature.
- All procedures requiring direct exposure of cells to air must be performed within a biosafety cabinet to ensure sterility and prevent contamination of the culture.
- Cells and NTOs are kept out of the incubator for as limited time as possible and all required steps need to be taken in a careful and timely manner.
- Only good quality ESCs (Figure 2A-C) form good quality NTOs (Figure 4B).
- If more than 10% of mouse ESCs are differentiated (Figure 2D), the culture is discarded and freshly thawed culture is prepared which is passaged at least twice before seeding ESCs for a differentiation assay (experienced researchers can judge by visual assessment (see also Mulas et al. ^21^) or via antibody staining for marker expression (e.g. OCT4, see also Kalkan et al. ^22^)).
- Cells and NTOs are kept without medium as briefly as possible to avoid drying out the culture and/or Matrigel.
- The optimal seeding density is recommended (Table 2 and Figure 3). However, for certain applications, such as early analysis, a seeding density up to 10 times higher may be advantageous.
- If NTOs look suboptimal (Figure 4C-E) or do not show correct spatiotemporal marker expression (Figures 5 & 6), the material does not meet quality standards and is discarded.

### 1. Maintenance passaging and seeding mouse ESCs for NTO differentiation

#### 1.1. Passaging

NOTE:

- ESCs are maintained on cell-culture dishes or wells of a 6-well plate pre-coated with 0.15% gelatin for 30 minutes at 37 °C.
- ESCs are passaged every 2-3 days, with the medium being replaced daily from day 2 onwards.
**1.1.1.** Check cells under phase-contrast microscope (see representative images, Figure 2).

**Figure 1:**
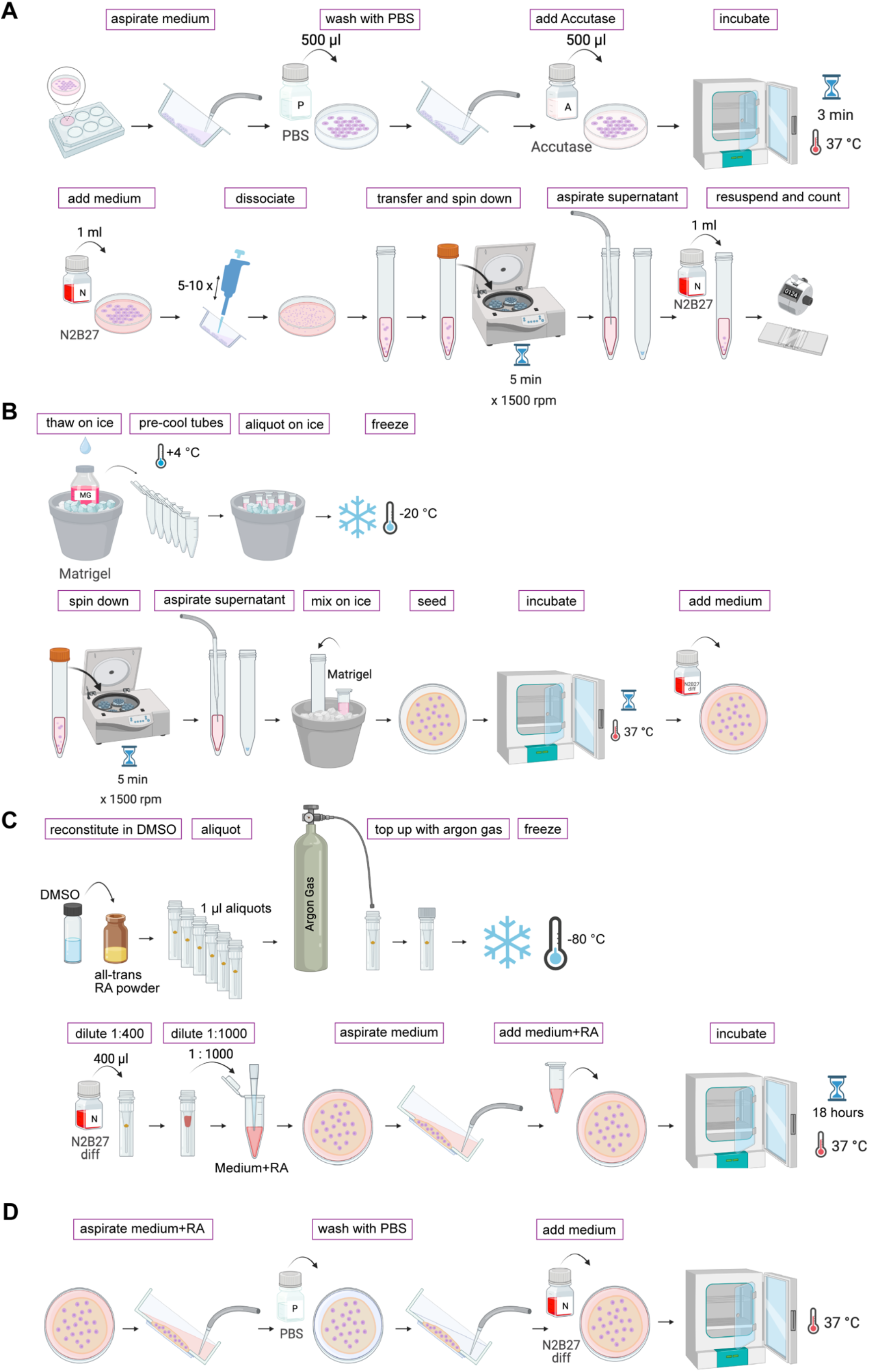
Overview of the protocol. **A:** Passaging ESCs (see **1.1.**). **B:** Aliquoting Matrigel and seeding single ESCs for differentiation into NTOs (see **1.2.**). **C:** Aliquoting RA and RA treatment (see **2.**). **D:** RA removal and routine of daily medium replacement (see **3.**). This figure was created with BioRender.com

**Figure 2:**
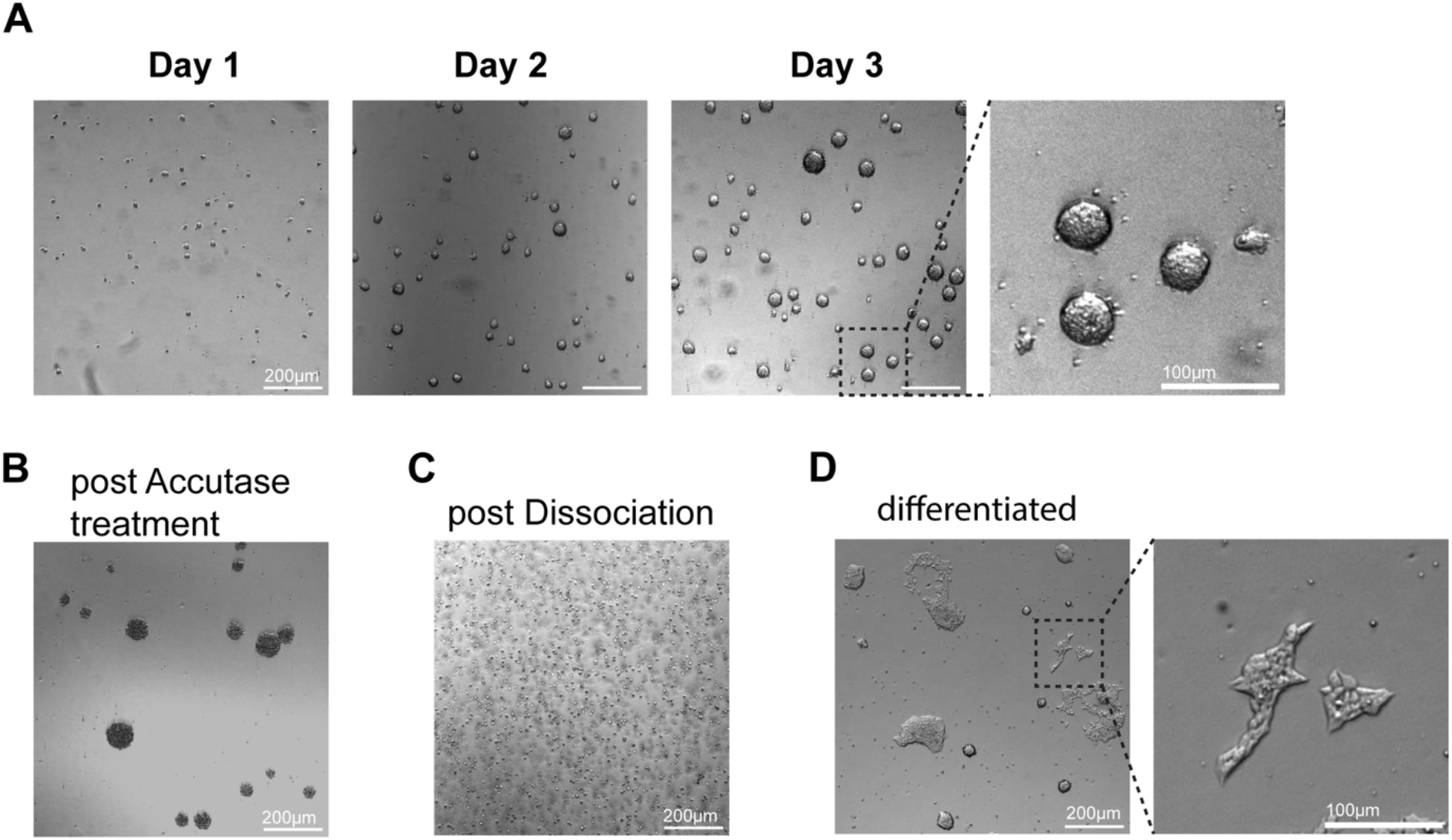
Good quality mouse ESCs (R1) maintained in N2B27 supplemented with 2iLIF (‘2iLIF’ in this protocol) **A:** Brightfield images of mouse ESCs 1-3 days after passaging. Colonies are doom- shaped and have a round, smooth outline. Scale bars, 200 µm (4x image) and 100 µm (zoom) **B:** Day 3 ESCs after 3 minutes of Accutase treatment at 37 °C. Colonies are detached from the bottom of the culture dish and single cells can be observed within the colony. Scale bar, 200 µm **C:** Colonies from B after dissociation into single cells. Scale bar, 200 µm **D:** Differentiated ESCs with colonies looking jagged (spikey edges) and flat. Scale bars, 200 µm (4x image) and 100 µm (zoom)

**Figure 3:**
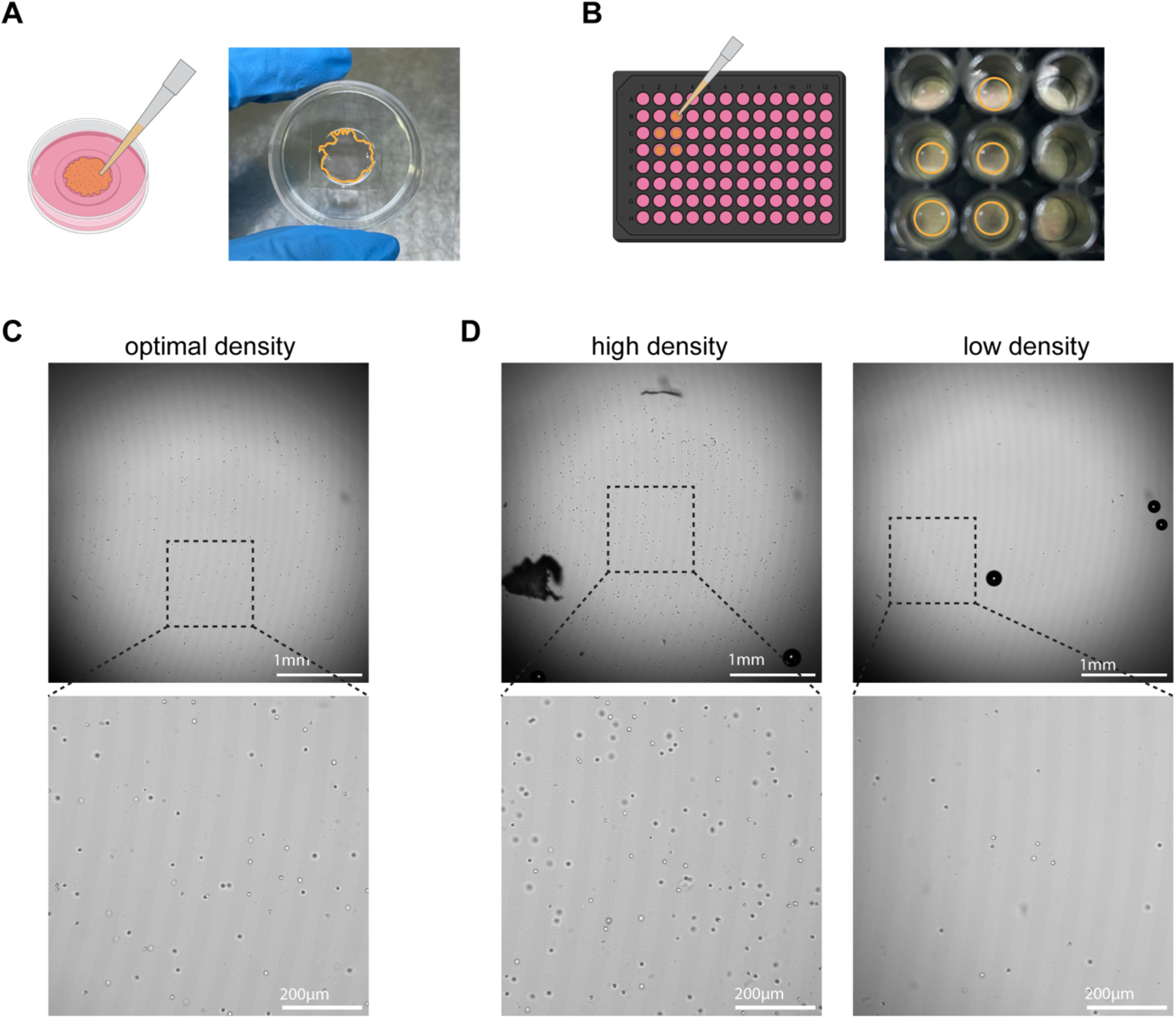
Seeding density in Matrigel in MatTek dish (A) and 96-well plate (B) **A:** Schematic of the Matrigel-cell suspension seeding process (orange) in a 35 mm glass bottom MatTek dish. Suspension is seeded using ‘flower’ technique to avoid majority of the cells being seeded close to the walls. **B:** Schematic of the Matrigel-cell suspension seeding process (orange) in a well of a 96-well plate. Suspension is seeded in a circle to avoid majority of the cells being seeded close to the walls. **C:** Optimal seeding density of 100 cells/ µl Matrigel (Table 2). Scale bars, 1 mm (4x image) and 200 µm (zoom) **D:** Suboptimal seeding density. Left panel: cell density too high (1000 cells/ µl Matrigel), right panel: cell density too low (10 cells/ µl Matrigel). Scale bars, 1 mm (4x image) and 200 µm (zoom)

**Figure 4:**
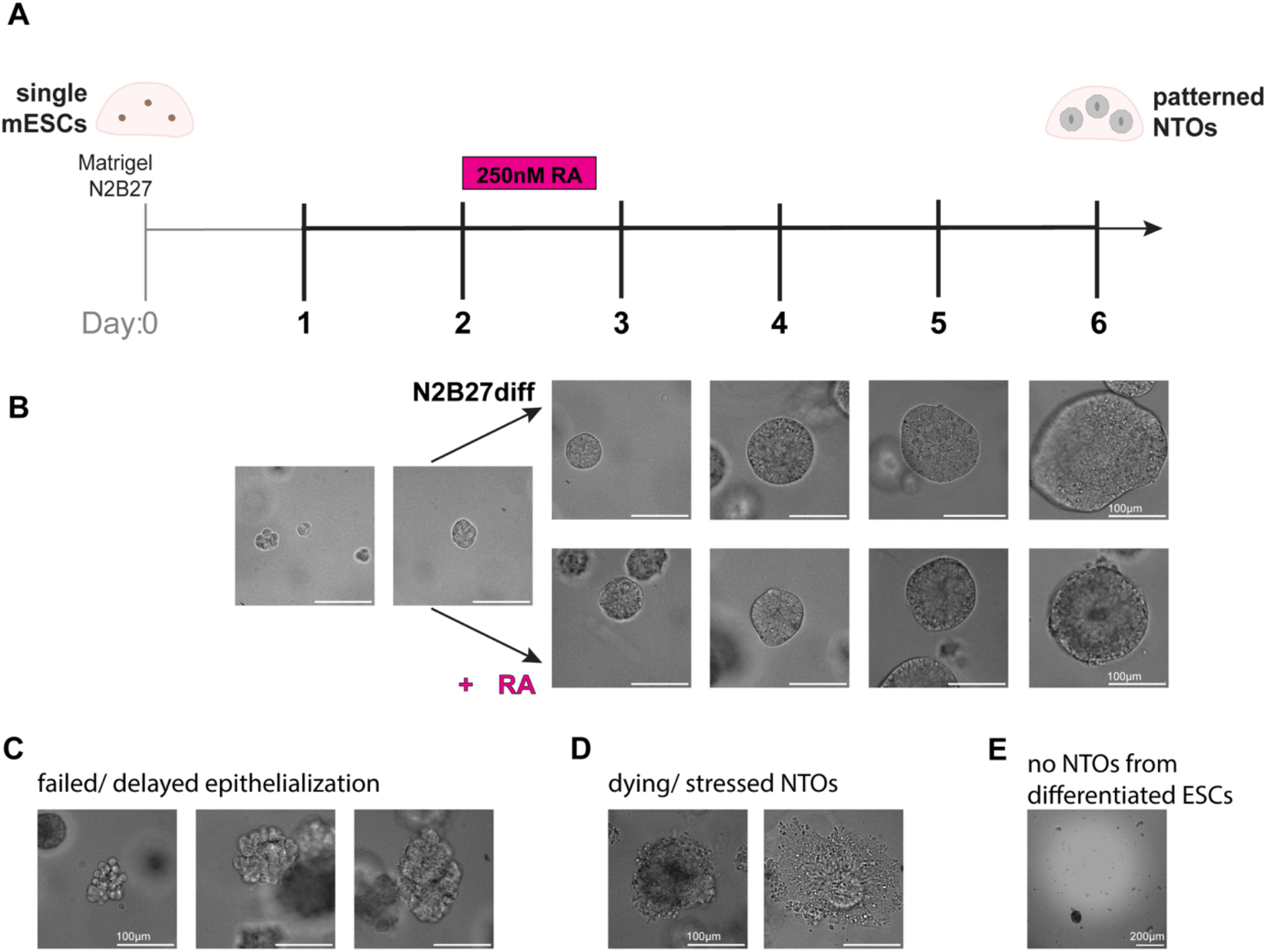
representative NTOs. **A:** Schematic of NTO differentiation protocol with 18 hours 250 nM RA pulse from day 2. **B:** Representative results of NTOs from day 1-6 according to timeline in A in N2B27diff (upper panel) or + RA (lower panel). Scale bars, 100 µm **C:** Example NTOs showing failed or delayed epithelialization. Scale bars, 100 µm **D:** Example NTOs showing dying or stressed NTOs. Scale bars, 100 µm **E:** Representation of day 6 ‘NTOs’ from differentiated ESCs. Scale bar, 200 µm

**Figure 5:**
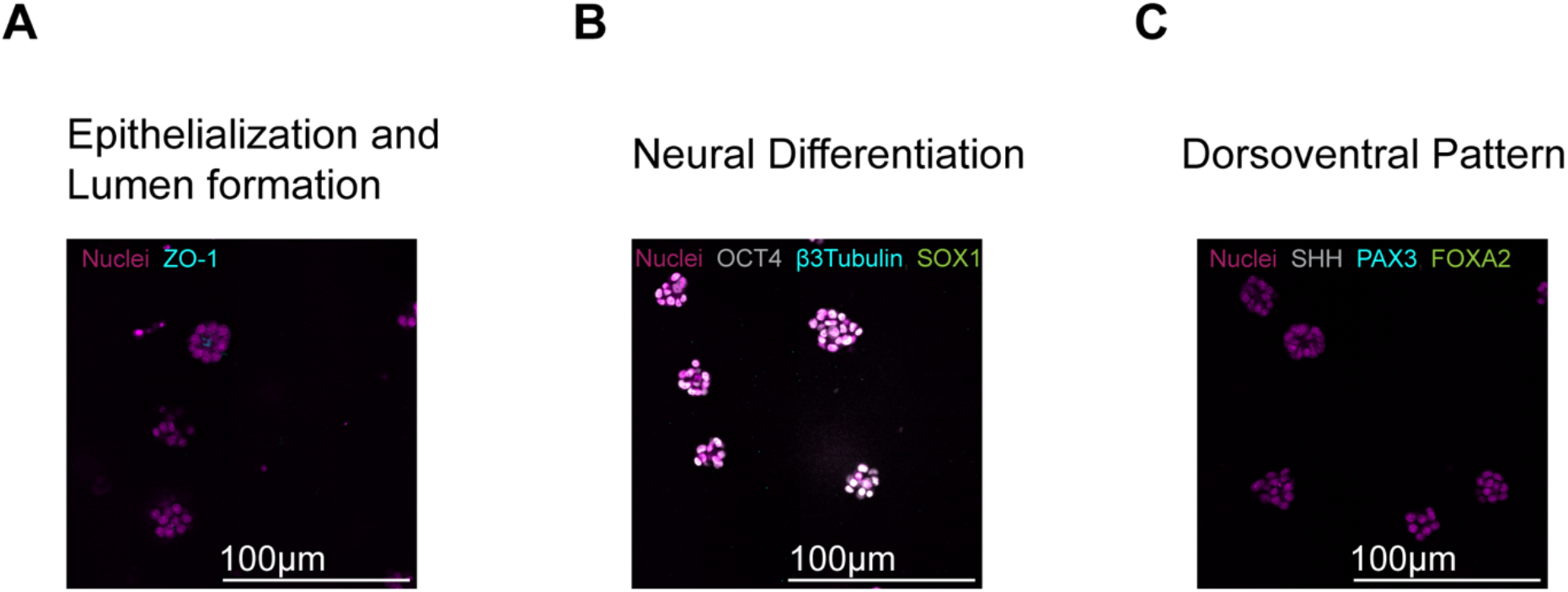
Marker expression in day 2 NTOs before RA treatment. **A:** Representative NTOs for epithelialization and lumen formation, with nuclei (magenta) and ZO-1 (cyan). Scale bars, 100 µm **B:** Representative NTOs for neural differentiation, with nuclei (magenta), OCT4 (grey), β3Tubulin (cyan) and SOX1 (green). Scale bars, 100 µm **C:** Representative NTOs for dorsoventral pattern, with nuclei (magenta), SHH (grey), PAX3 (cyan) and FOXA2 (green). Scale bars, 100 µm

**Table 1:** Medium components (N2B27 base, 2iLIF, N2B27diff)

| N2B27 base (for 50 ml) |  | 2iLIF (for 50 ml) |  | N2B27 diff (for 50 ml) |  |
| --- | --- | --- | --- | --- | --- |
| 25 ml | Neurobasal | 50 ml | N2B27 base | 50 ml | N2B27 base |
| 25 ml | DMEM/ F12 | 20 ng/ml | mouse LIF | 0.5 ml | Sodium Pyruvate |
| 0.5 ml | B27 | 3 μM | CHIR 99021 | | |
| 0.25 ml | N2 | 1 μM | PD 0325901 | | |
| 0.5 ml | Pen/ Strep |  |  |  |  |
| 0.5 ml | Glutamine |  |  |  |  |
| 0.1 mM | β-Mercaptoethanol | | | | |

**Table 2:** Information on cell number per dish, amount of Matrigel, volume of medium and jellification time.

| Dish/ well | Matrigel | Medium/ Volume | Cells (at 100 cells/ μl Matrigel density) | Jellification time (37 °C) |
| --- | --- | --- | --- | --- |
| MatTek (35mm glass bottom dish) | 40 μl | 2 ml | 4000 | 15 minutes |
| 96-well plate | 4 μl | 100 – 200 μl (depending on dish) | 400 | 4 minutes |
| 24-well plate | 10 – 15 μl | 1 ml | 1000 – 1500 | 10 minutes |
| 6-well plate | 40 – 80 μl | 2 ml | 4000 – 8000 | 15 minutes |

NOTE:

- Ensure that colonies have an appropriate size and proper morphology (Figure 2A), while also confirming that they adhere to the bottom of the dish.

**1.1.2.** Open the lid of the culture dish/ plate, slightly tilt it and carefully aspirate medium.
**1.1.3.** Wash colonies with 500 µl pre-warmed PBS (added to cells in a single well of a 6-well plate), add and then aspirate again immediately.

NOTE:

- This step is crucial because: A) ESC medium (2iLIF) carry over must be avoided prior to the differentiation assay (or NTO formation will be hindered), B) FBS or BSA in culture medium can stop dissociation. This applies to ESCs cultured in a different ESC medium, containing FBS or BSA.

**1.1.4.** Add 500 µl pre-warmed Accutase and transfer cells back into the incubator.
**1.1.5.** Incubate cells at 37 °C for 3 minutes using a timer.
**1.1.6.** After the incubation time, take out the plate and check cells under the microscope.

NOTE:

- ESCs should have detached from the culture plate, if not tap the dish/plate firmly but carefully (Figure 2B).

**1.1.7.** Add 1 ml N2B27 medium by pipetting it directly onto to cells.

NOTE:

- ESCs should be washed off the plate, leaving ‘empty’ areas in the culture dish.

**1.1.8.** Take a P1000 pipette, tilt the plate slightly such that all the colonies sticking to the culture dish can be washed off.
**1.1.9.** To dissociate clumps of cells, tilt the plate slightly, place the pipette tip (P1000) in the middle of the dish. Press the tip against the bottom of the dish but slightly off the perpendicular axis and flush out the cells. Repeat for five to ten times to break the cell clumps.
**1.1.10.** Check that the cells are dissociated properly into single cells under the microscope (Figure 2C), repeat step 1.1.9. if there are still clumps.
**1.1.11.** After complete dissociation, transfer the cell suspension into a clean sterile 15 ml Falcon tube, wash the empty well where cells were cultured in with 1 ml N2B27 medium and transfer to the same Falcon tube with the cell suspension.
**1.1.12.** Spin down cells for 5 minutes at 1500 rpm in an ultra centrifuge.
**1.1.13.** Carefully aspirate the supernatant.
**1.1.14.** Resuspend the cells in an appropriate amount of N2B27 medium (e.g. 1 ml) and count cells (see Table 2 for recommended cell numbers for differentiation assay into NTOs).

#### 1.2. Differentiation of ESCs into neural tube organoids (NTOs)

##### 1.2.1. Aliquoting Matrigel

**1.2.1.1.** Thaw bottle of Matrigel overnight on ice at 4 °C.

NOTE:

- Best to do this in a box with ice at 4 °C or in the fridge
- Do not leave Matrigel for longer than necessary at 4 °C, aliquot and freeze as soon as possible (otherwise this can affect the functionality of the Matrigel)

**1.2.1.2.** In the biosafety cabinet: once the Matrigel is thawed, place 1.5-2 ml Eppendorf tubes on ice to pre-cool them, while also keeping the Matrigel bottle on fresh ice.
**1.2.1.3.** Mix the thawed Matrigel carefully to avoid bubble formation.
**1.2.1.4.** Aliquot Matrigel at the desired volume (0.5-1 ml is recommended) and freeze at −20 °C until further usage.

NOTE:

- Do not replace the tip between the aliquots, these are ‘warm’ (room temperature) and will cause the Matrigel to jellify within, which leads to a significant loss of Matrigel.
- Instead, cool the pipette tips to 4 °C by putting them in the fridge before use.

##### 1.2.2. Seeding single cells in Matrigel

**1.2.2.1.** Quantify the cell number needed (100 cells/ µl Matrigel, see Table 2).
**1.2.2.2.** Transfer appropriate cell number (cell suspension) to a clean 15ml Falcon tube and top it up with 1ml of N2B27.
**1.2.2.3.** Spin down cells for 5 minutes at 1500 rpm in the centrifuge.
**1.2.2.4.** Aspirate the supernatant and place Falcon tube on ice.
**1.2.2.5.** Carefully resuspend the cell pellet on ice in an appropriate amount of thawed and liquid Matrigel while avoiding the formation of bubbles.
**1.2.2.6.** Spread the appropriate amount of Matrigel-cell suspension, in a thin layer, onto a glass bottom or plastic bottom dish which is optimal for imaging (see Table 2, Figure 3A-C).

NOTE:

- Make sure not to touch the walls of multi-well plates.
- If using multi-well plates: best to first draw a circle of Matrigel using a P20 (96-well plate) or P200 pipette tip before ‘filling it’ with the Matrigel-cell suspension (Figure 3B).

**1.2.2.7.** Put the multi-well plate/ dish in the incubator at 37°C to let the Matrigel jellify (rule of thumb: 1 minute per µl Matrigel for smaller amounts and a maximum of 15 minutes for 40 µl or more) (use a timer).
**1.2.2.8.** After the incubation time, take out the plate and add appropriate volume of medium (see Table 2).
**1.2.2.9.** Check the seeding density and cell distribution within the Matrigel under the microscope (Figure 3C-D).

NOTE:

- Keep Matrigel aliquot on ice or in the fridge, otherwise it will jellify at room temperature.
- Carefully mix the Matrigel-cell suspension by resuspending with a P300 or P1000 pipette tip before seeding for differentiation assay, otherwise cell number will be highly variable between wells/ dishes (not recommended to do before seeding every single well, because this leads to loss of Matrigel as Matrigel can jellify and stick to the pipette tip).
- The presence of bubbles into the Matrigel can cause its detachment off the well/ dish after jellification.
- ESCs in Matrigel should not dry out during seeding for differentiation assay into NTOs or replacing medium. After jellification, medium should be applied as soon as possible. Jellification typically is complete in 1 minute at 37 °C per 1 µl Matrigel and maximum 15 minutes for 40 – 100 µl Matrigel when spread as a thin layer.
- Because of high surface tension, some multi-well plates can cause the Matrigel droplet to move from the center of the well to the walls if the well walls are touched, which causes an inadequate distribution of cells/ NTOs.
- Matrigel should always be thawed on ice or 4 °C (never at room temperature or in hands).

### 2. RA treatment of NTOs to enhance neural differentiation and to induce posteriorization and ventralization

**2.1.** Aliquoting retinoic acid (RA)

**2.1.1.** Bring all-trans RA powder to room temperature (stored at −20 °C).
**2.1.2.** Reconstitute powder in an appropriate volume of DMSO to reach a final concentration of 100 mM.
**2.1.3.** Aliquot 1 µl of 100 mM RA into one 1.5 ml screw-top vial each.
**2.1.4.** Add Argon gas until tube is ‘full’.

NOTE:

- Since argon gas is heavier than oxygen, it will sink to the bottom of the tube to cover and preserve RA from air. This can be felt on hands when it leaks out when tube is full.

**2.1.5.** Tightly close the vial.
**2.1.6.** Store aliquots in a black box at −80 °C until further usage (alternatively: aliquot in Eppendorf tubes and store in a black box in liquid N_2_).

NOTE:

- RA is highly sensitive to light and air.
- Keep thawed RA aliquots always in the dark.
- Reconstituted and diluted RA can be stored in the fridge (covered with foil) for up to 2 days (after that, discard and use a fresh aliquot).

**2.2.** RA treatment of NTOs for 18h

**2.2.1.** Take out 1 µl aliquot with a concentration of 100 mM from −80 °C and keep dark until thawed on room temperature (palm of hand or in tube rack in biosafety cabinet without lights on).

NOTE:

- If RA aliquots are stored in liquid N_2_, immediately open the lid of the Eppendorf tube to avoid explosion of the tube, then close again and proceed to step 2.2.2.

**2.2.2.** Once thawed, add 400 µl of pre-warmed N2B27diff medium for a concentration of 250 µM using a P1000 pipette. Mix thoroughly to achieve homogeneity.

NOTE:

- Alternatively, RA can be reconstituted and diluted in pure DMSO instead of N2B27diff medium, although this increases the concentration of DMSO in the culture.

**2.2.3.** Use the dilution from 2.2.2. in a ratio of 1:1000 in N2B27diff to get an appropriate volume with a final concentration of 250 nM RA.
**2.2.4.** Take out plates/ dishes growing NTOs from the incubator and place into a sterile biosafety cabinet.
**2.2.5.** Remove the lid of the plate/ dish and slightly tilt the plate to the side.
**2.2.6.** Aspirate the medium carefully either using a P200 pipette, or an aspirator without disturbing the Matrigel.
**2.2.7.** Add an appropriate volume of N2B27diff supplemented with 250 nM RA solution (see Table 2), cover with the lid and transfer the NTOs back into the incubator.

NOTE:

- Sometimes, especially if the RA was not mixed completely, precipitates can form that are harder to dissolve in normal N2B27diff (then, use DMSO). In this case it is advisable to discard the aliquot and make fresh RA aliquots because these precipitates have an unknown and uncontrolled concentration of RA.
- Make sure ESCs in Matrigel do not dry out while replacing medium or treating with RA (handle a maximum of 8 wells of 96-well plate if using a multichannel pipette, 1 MatTek dish/ well of any other multi-well plate)

### 3. RA removal and routine of daily medium replacement of NTOs

**3.1.** Transfer the culture plate from the incubator and move to the sterile biosafety cabinet.
**3.2.** Remove the lid and slightly tilt the plate to the side.
**3.3.** Carefully aspirate N2B27diff containing RA either using a P200 pipette, or aspirator without touching the Matrigel.
**3.4.** Add pre-warmed PBS (add enough volume to cover the Matrigel droplet, or use volume as referred to in Table 2), then tilt the plate again and carefully remove it to wash out the leftover RA.
**3.5.** Add appropriate volume of N2B27diff by pipetting the medium at the wall of the well/ dish to not destroy the Matrigel droplet.
**3.6.** Replace medium daily following the steps 3.1. – 3.3., then adding fresh pre-warmed N2B27diff by pipetting the medium onto the wall of the well/ dish before returning the NTOs back in the incubator.

NOTE:

- Direct and vigorous pipetting directly or addition of cold medium/ PBS onto the Matrigel droplet can cause its breakage and detachment from the plate/dish.
- When perturbing pathways by using small molecules or recombinant proteins, it is important to replace the medium daily. If drugs are to be applied for longer than one day, drug solutions need to be prepared freshly daily or at least every two days to ensure protein/ drug stability and the continued presence of correct concentrations.

### 4. Fixation of NTOs

NOTE:

- NTOs are fixed as described by Krammer et al.^19^. Depending on the antibody to be subsequently used, PFA from different sources (store bought, in-house, etc.) can be required.

**4.1.** Thaw PFA aliquot at 4 °C overnight.
**4.2.** Bring PFA to room temperature before use.
**4.3.** If using a higher stock concentration of PFA, dilute PFA to 4% in N2B27diff medium or PBS in a chemical hood.
**4.4.** Take NTOs out of the incubator and transfer them to a chemical hood.
**4.5.** Open the lid and add 4% PFA in a ratio of 1:1 directly to the NTOs to result in a final concentration of 2% (e.g. 100 µl 4% PFA added to the 100 µl N2B27diff in the 96-well plate).
**4.6.** Put the lid back on and incubate NTOs for 30 minutes at room temperature (set a timer).

NOTE:

- PFA is toxic and should not be inhaled. Work carefully and with caution under the chemical hood.
- Longer fixation can impair performance of certain antibody treatment.
- Some antibodies require usage of a specific PFA or shorter/ optimized fixation time.

**4.7.** After 30 minutes, remove the lid and slightly tilt the plate to the side, then follow steps 3.3. – 3.4.
**4.8.** Carefully add pre-warmed PBS (add enough volume to cover the Matrigel droplet or use volume as referred to in Table 2), and incubate for 5 minutes at room temperature, repeat this step 3 times to remove all left over PFA.
**4.9.** After the final wash, add appropriate amount of PBS (see Table 2), seal the plate using parafilm and store at 4 °C until further usage.

### 5. Antibody staining of NTOs

NOTE:

- Antibody staining is a valuable in assessing the correct identity of the NTOs as well as the success of the assay/ experiment.
- Antibody staining can be performed as outlined in Meinhardt et al. ^17^ and Krammer et al. ^19^
- Block/ permeabilization solution (1x PBS with 1% BSA and 0.5% Triton-X) is kept at 4 °C and adapted to room temperature before use.
- For a full 3D reconstitution and analysis after imaging, antibodies (1^st^ and 2^nd^) should be kept at least 1, but preferably 2 – 3 nights at 4 °C to ensure good antibody penetration.
- Alexa Fluor anti-561 2^nd^ antibody might give stronger background signal in Matrigel in the first three days of the differentiation assay.

### 6. Clearing of NTOs

NOTE:

- NTOs can be cleared using the protocol from Krammer et al.^19^. The addition of propylgallate to the clearing solution minimizes photobleaching of the NTOs.
- Clear NTOs only after completing the antibody staining process (1^st^ and 2^nd^ antibodies are incubated and washed out properly).
- Perform clearing approximately 1-2 hours before imaging to prevent oxidization of the clearing solution (liquid turns from light yellow/cream into dark bright yellow or even brown) or crystallization in the well.
- Crystallized clearing solution can be dissolved by the addition of PBS.
- Correct pH of the clearing solution is crucial, otherwise Matrigel can dissolve or clearing solution can oxidize at a faster rate.
- Avoid shaking or mixing the clearing solution harshly because this results in the formation of air bubbles.

### 7. Analysis of NTOs

To analyze the NTOs after full penetration of antibody staining and whole tissue clearing, we recommend using a confocal microscope with 10-40x objectives. For 3D analysis, the software CellProfiler ^23, 24^, Imaris or similar can be used. Verifying the correct identity of NTOs and the spatiotemporal expression of specific dorsoventral patterning markers is essential to determine whether the differentiation assay has been successful before addressing fundamental research questions. In Figures 5 and 6, we show characterization of three essential aspects to determine the success of the culture:

1. Epithelialization and lumen formation (ZO-1)
2. Neural differentiation (OCT4, SOX1, β3-tubulin) (in-depth characterization of neural differentiation trajectory in ^17^)
3. Dorsoventral patterning (FOXA2, SHH, PAX3) (in-depth characterization of ventral patterning in ^19^)

**Figure 6:**
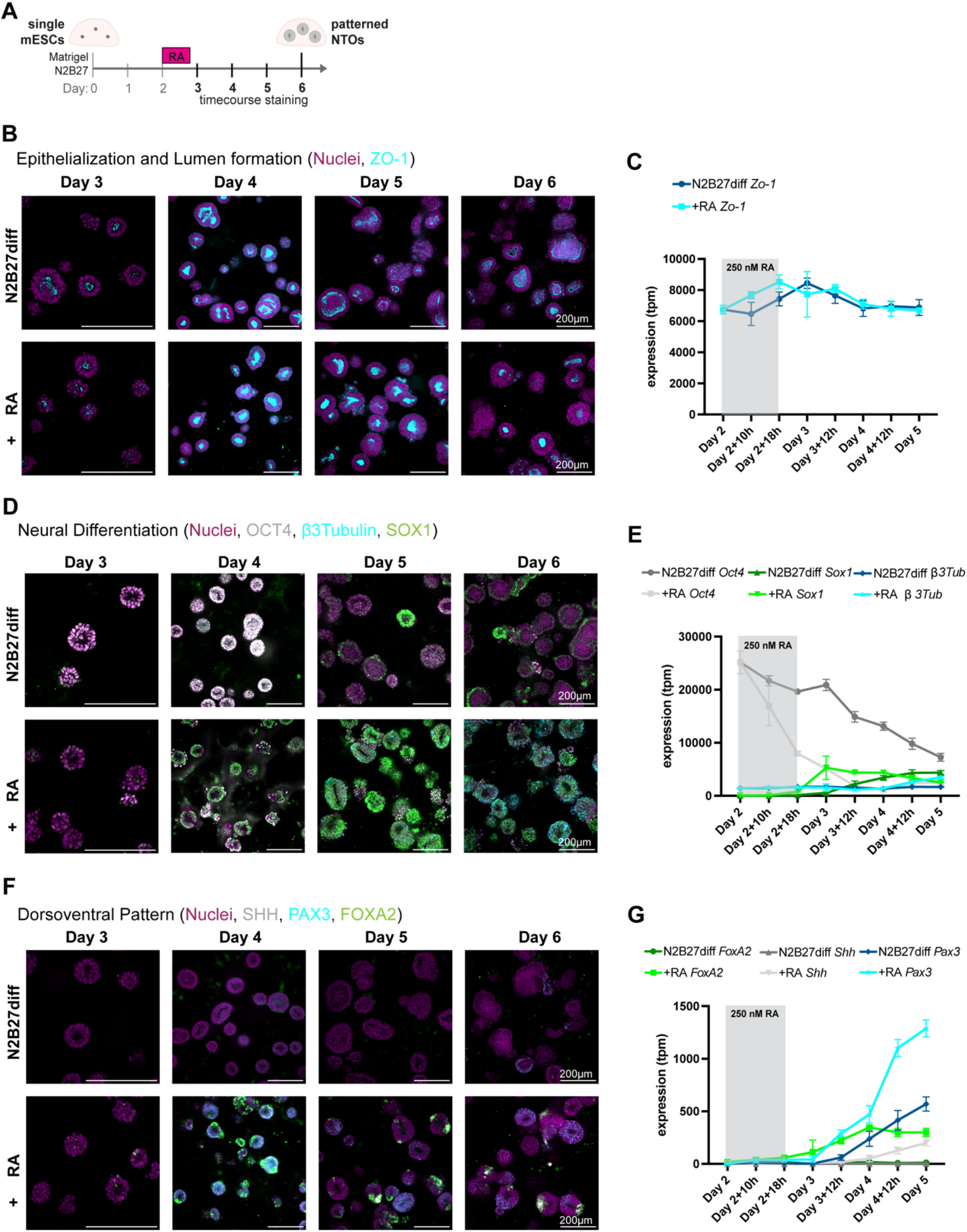
Marker expression from day 3-6 in control (N2B27diff) and RA-treated (+ RA) NTOs. **A:** Schematic of the NTO differentiation assay with 18 hours 250 nM RA pulse from day 2 and timecourse staining from day 3-6. **B:** Representative timecourse of NTOs for epithelialization and lumen formation, with nuclei (magenta) and ZO-1 (cyan). Scale bars, 200 µm **C:** Timecourse RNA sequencing^25^ for *Zo-1* (dark blue respectively cyan) from day 2-5 in N2B27diff and + RA. **D:** Representative timecourse of NTOs for neural differentiation, with nuclei (magenta), OCT4 (grey), β3Tubulin (cyan) and SOX1 (green). Scale bars, 100 µm **E:** Timecourse RNA sequencing^25^ for *Oct4* (dark respectively light grey), *Sox1* (dark respectively light green) and *β3Tubulin* (dark blue respectively cyan) from day 2-5 in N2B27diff and + RA. **F:** Representative timecourse of NTOs for dorsoventral pattern, with nuclei (magenta), SHH (grey), PAX3 (cyan) and FOXA2 (green). Scale bars, 200 µm **G:** Timecourse RNA sequencing^25^ for *Shh* (dark respectively light grey)*, Pax3* (dark blue respectively cyan) and *FoxA2* (dark respectively light green) from day 2-5 in N2B27diff and + RA.

## REPRESENTATIVE RESULTS

R1 mouse ESCs were used for all representative examples in figures presented in this manuscript. Other mentioned mouse ESCs can exhibit mild differences in epithelialization timing, neural differentiation or patterning efficiency when seeded for NTOs, however, results should not significantly differ from the data shown here.

When good quality ESCs (Figure 2) are seeded at the appropriate density (Figure 3) in Matrigel, these single cells clonally expand and epithelialize within two days (Figure 4). NTOs exhibit a smooth, round-to-ellipsoid shape with a single lumen. NTOs that have failed to epithelialize lack lumen formation, with individual cells remaining visible. Small round debris observed around the NTOs are apoptotic bodies, indicative of a stressed culture.

On day 2, NTOs are negative for any neural (SOX1) and neuronal (β3 Tubulin) marker, as well as dorsoventral patterning markers (FOXA2, PAX3, SHH), but are expressing ZO-1 and OCT4 (Figure 5). On day 2, a 250 nM RA pulse is applied for 18 hours to ventralize the NTOs to hindbrain/ cervical spinal cord levels and to boost neural differentiation ^17, 19^. From day 2 onwards, NTOs gradually downregulate the pluripotency marker OCT4 and from day 3 onwards, they start expressing SOX1 and subsequently β3 Tubulin by day 6 (Figure 6). RA treatment accelerates neural differentiation compared to NTOs in N2B27diff only. Full dorsoventral patterning markers (FOXA2, SHH, PAX3) are only expressed after RA treatment, while without RA, NTOs will become positive for PAX3 and SOX1 by day 6 (Figure 6). RA-treated NTOs exhibit a smaller and more compact lumen compared to those in N2B27diff, as demonstrated by the tight-junction marker ZO-1 (Figures 4 and 6).

## DISCUSSION

Here, we present several optimizing adaptations and improvements to the original protocol for generating high quality NTOs, to facilitate standardization across laboratories. Key changes include different ESC culture conditions, reduced input cell numbers, improved differentiation medium, modified differentiation protocol, and seeding density adjustments for high-throughput analysis using multi-well plates.

The most critical points of the protocol are:

- Source and use of the correct Matrigel with appropriate protein concentration (inappropriate protein concentration can affect NTO patterning efficiency, see Table 3)
- Seeding of single cells in the correct density (seeding clumps of cells can lead to ‘grape’-formation and non-epithelialized NTOs with multiple lumen ^25^)
- Complete removal of 2iLIF from ESC maintenance medium before seeding for NTOs (carry-over of 2iLIF can impair NTO epithelialization, see Table 3)
- RA treatment using 250 nM for 18 hours on day 2 is crucial for pattern formation as tested in ^17^ (lower concentration and/or shorter pulse decreases pattern efficiency)
- Fixation with 2% PFA (using higher concentrated PFA can dissolve Matrigel)
- Prolonged staining combined with overnight antibody incubation to ensure full penetration that allows for subsequent 3D analysis (shorter incubation time can cause incomplete penetration and thus impair downstream analysis)
- Correct refractive index and pH of the clearing solution, and appropriate time of addition (only after antibody staining and shortly before imaging) (for structures >200µm 30-60 minutes prior to imaging are sufficient)

**Table 3:**
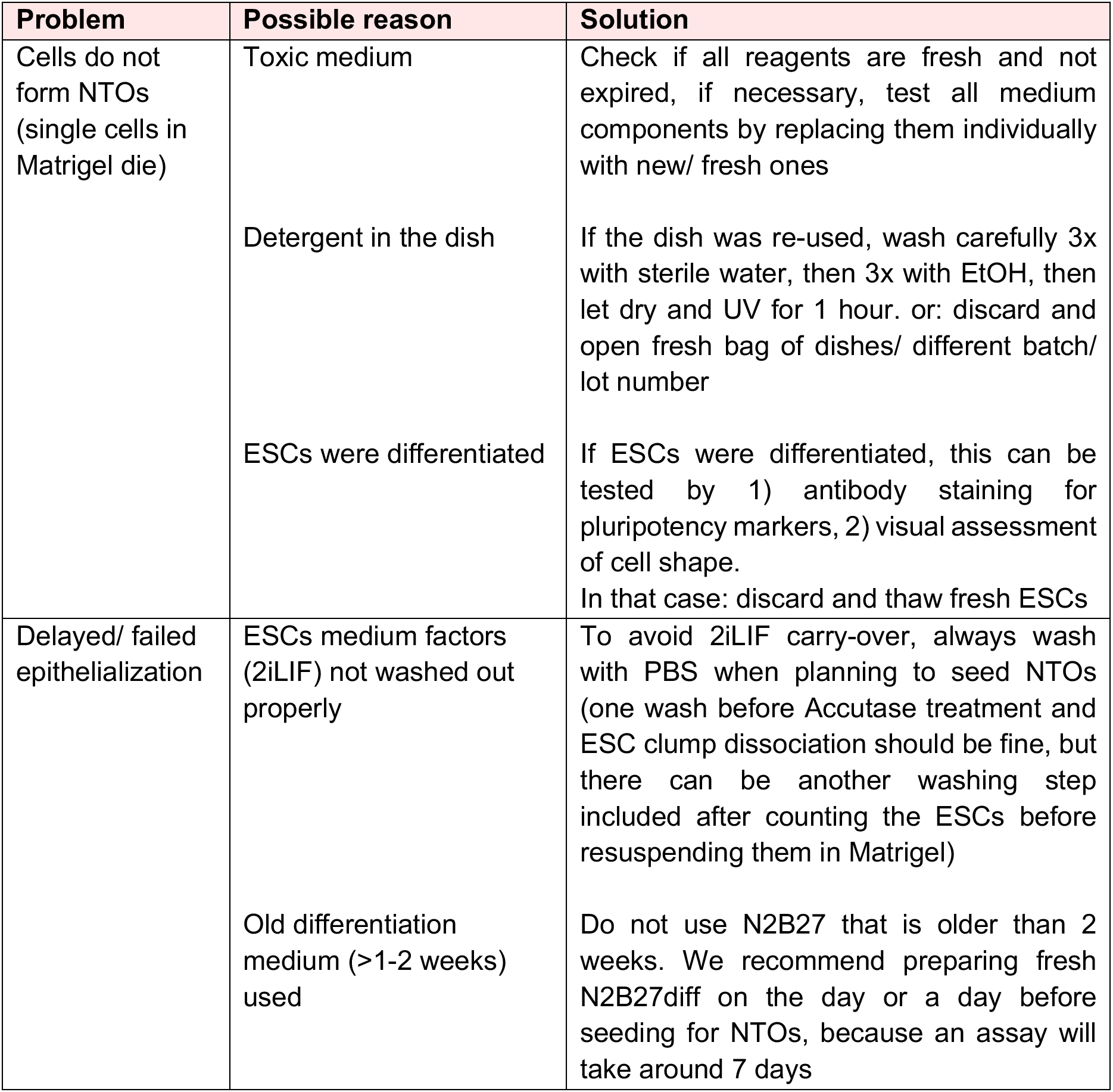
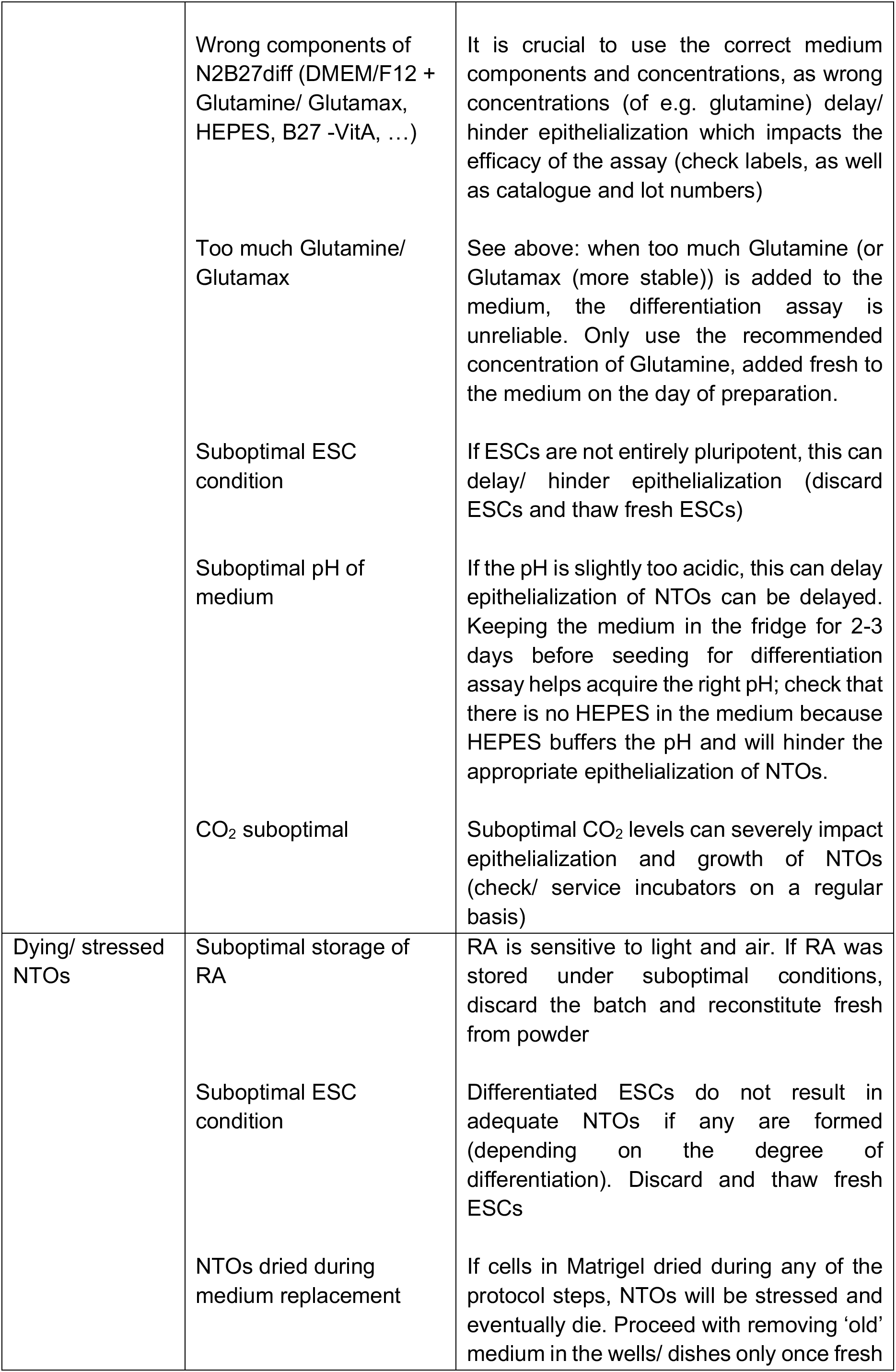
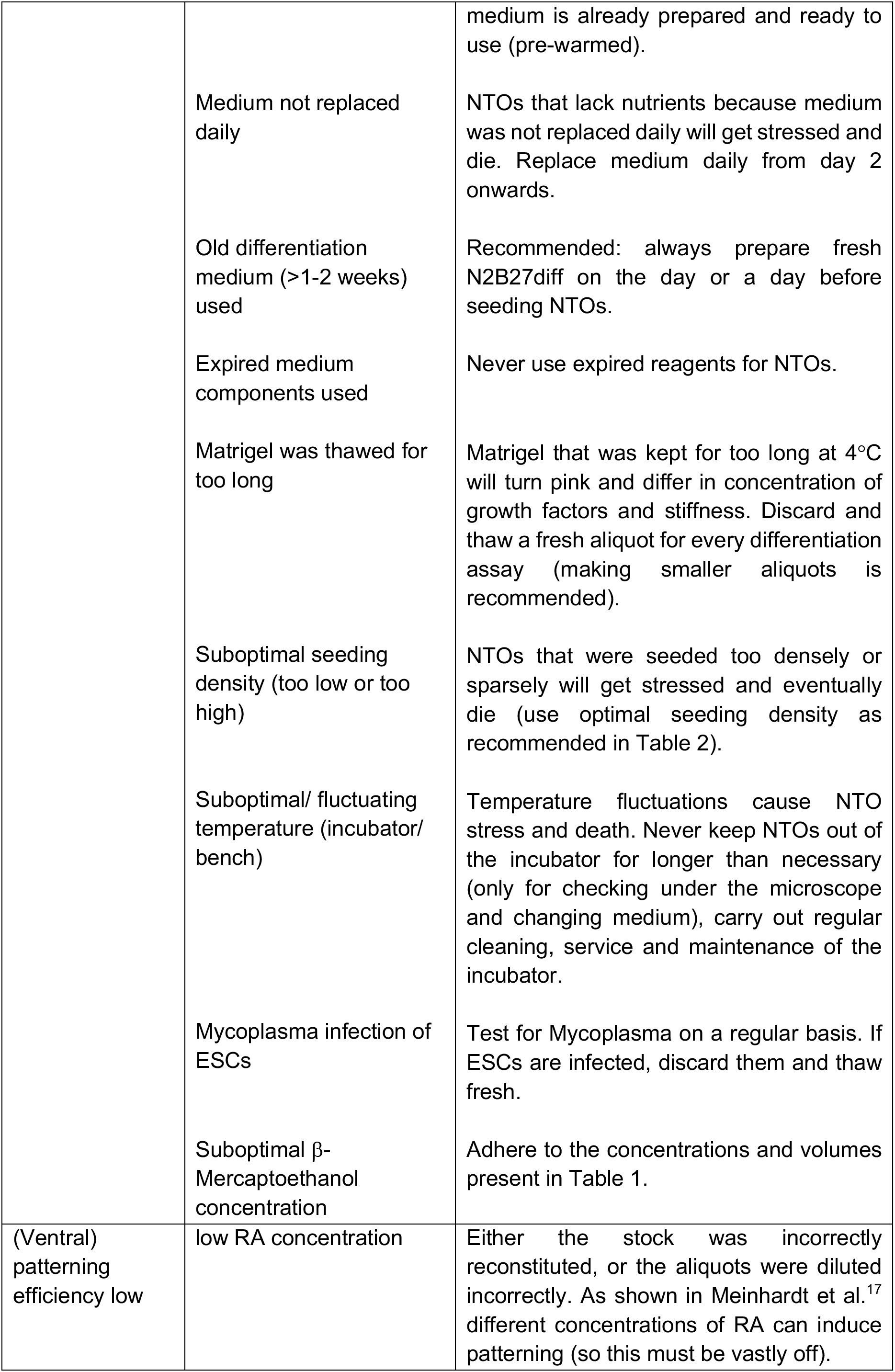
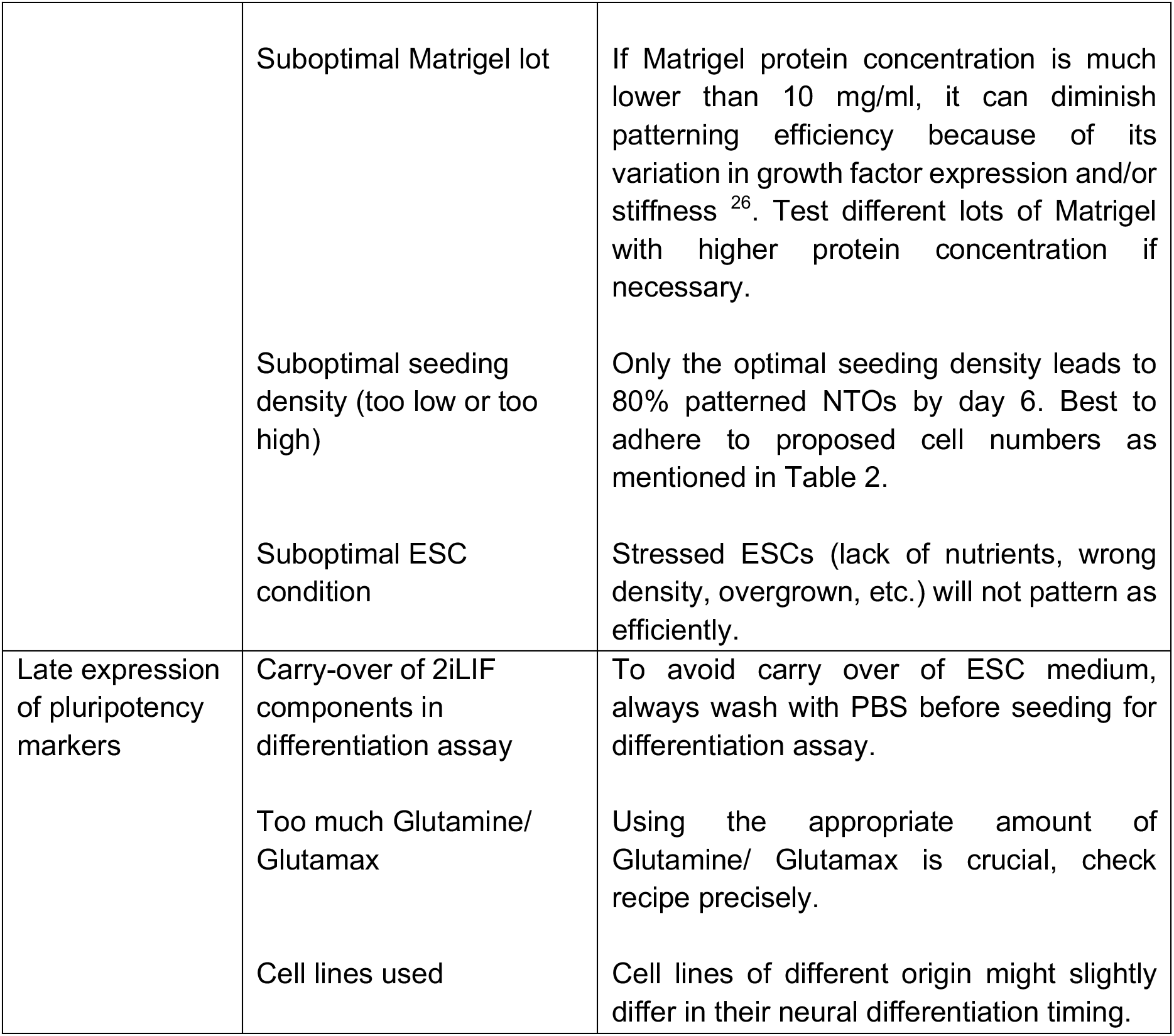
Troubleshooting of the method.

Furthermore, establishing a good quality and homogenously pluripotent ESC culture^21^ is crucial for the culture’s successful and reproducible differentiation into NTOs. Using ‘suboptimal’ quality cells leads to marked inconsistencies in data acquired, sacrificing time and resources. Differentiated ESCs will not form NTOs and die as single cells in Matrigel. The following points are critical and need to be checked precisely to culture good quality ESCs:

- Culture conditions: ideally, 50 000 cells are seeded per passage, in a well of a 6-well plate with passaging every 2-3 days and medium replacement daily from day 2. Inconsistency in seeding density and irregularity in medium replacement can have severe negative effects on the quality of the cells (see Table 3 for more details). We recommend to culture ESCs in 2iLIF medium on 0.15% gelatin-coated plates as this provides the most homogenously pluripotent starting culture ^21^.
- Differentiation: differentiated cells will not be forming NTOs but remain single cells in the Matrigel and die (which results in lower number of ‘good’ NTOs, see Table 3).
- Contamination: Mycoplasma contamination delays NTO formation and growth, resulting in smaller, stressed organoids that eventually die. Neural differentiation and patterning efficiency are also severely affected when ESCs contaminated with mycoplasma.

This protocol has been rigorously tested using the cell lines R1^27^, 46C^28, 29^, IB10^30^, ES-E14TG2a^31–33^, 129S6^34^, HM1^33^, and we confirmed that they feasibly made NTOs. Other cell lines (e.g. with C57BL/6 background), might require optimizations. Slight differences in anterior-posterior identity, dorsoventral patterning efficiency, and degree of ventralization can be observed among different cell lines. We acknowledge the presence of variability and heterogeneity in organoid size, despite starting from single cells. Because Matrigel is an expensive, animal-derived resource that is often sparsely available and has unpredictable delivery times, PEG-based hydrogels can be used alternatively ^26^.

This protocol was optimized from Meinhardt et al.^17, 20^ to increase survival of ESCs in Matrigel, as well as epithelialization and patterning efficiency of NTOs.

- Defined, more homogenous pluripotent ESC culture.
- Lower cell number required (due to homogeneous pluripotent starting culture, more single cells survive and form NTOs).
- Increased patterning efficiency (from approx. 40% (Meinhardt et al.) to 80%) due to factors mentioned above together with updated differentiation medium formulation (new in this protocol).
- No Noggin supplementation is required in the initial two days before the RA pulse.
- High throughput analysis possible due to adaption to multi-well usage.

To our knowledge, this neural organoid protocol uniquely enables progeny from a single cell to spontaneously undergo patterning upon self-organization following a global pulse of RA in a defined medium, without the need for additional factors or microfluidics to mimic spatial stimulus gradients.

NTOs can be used to study aspects of floorplate self-organization such as reported by Krammer et al. 2024 ^19^. This work demonstrates the formation of organizers, and it challenges the previous notion that embryonic induction relies solely on a controlling structure, such as the notochord. Instead, it reveals that these processes can occur through the regulated interconnections among cells.

Studying self-organization in an organoid system can give us novel insights into patterning in the developing organism. By manipulating various factors during NTO formation, epithelialization or patterning, key regulators of neural tube development, the underlying mechanisms of certain developmental disorders and potential targets for intervention can be identified. Furthermore, NTOs offer a platform for drug screening and development, where the efficacy and safety of potential therapeutic compounds can be tested in a more physiologically relevant environment.

This protocol is a standard in the Tanaka laboratory and highlights the versatility of NTOs and their potential to advance various areas of neuroscience, developmental biology, and regenerative medicine.

## TABLE OF MATERIALS

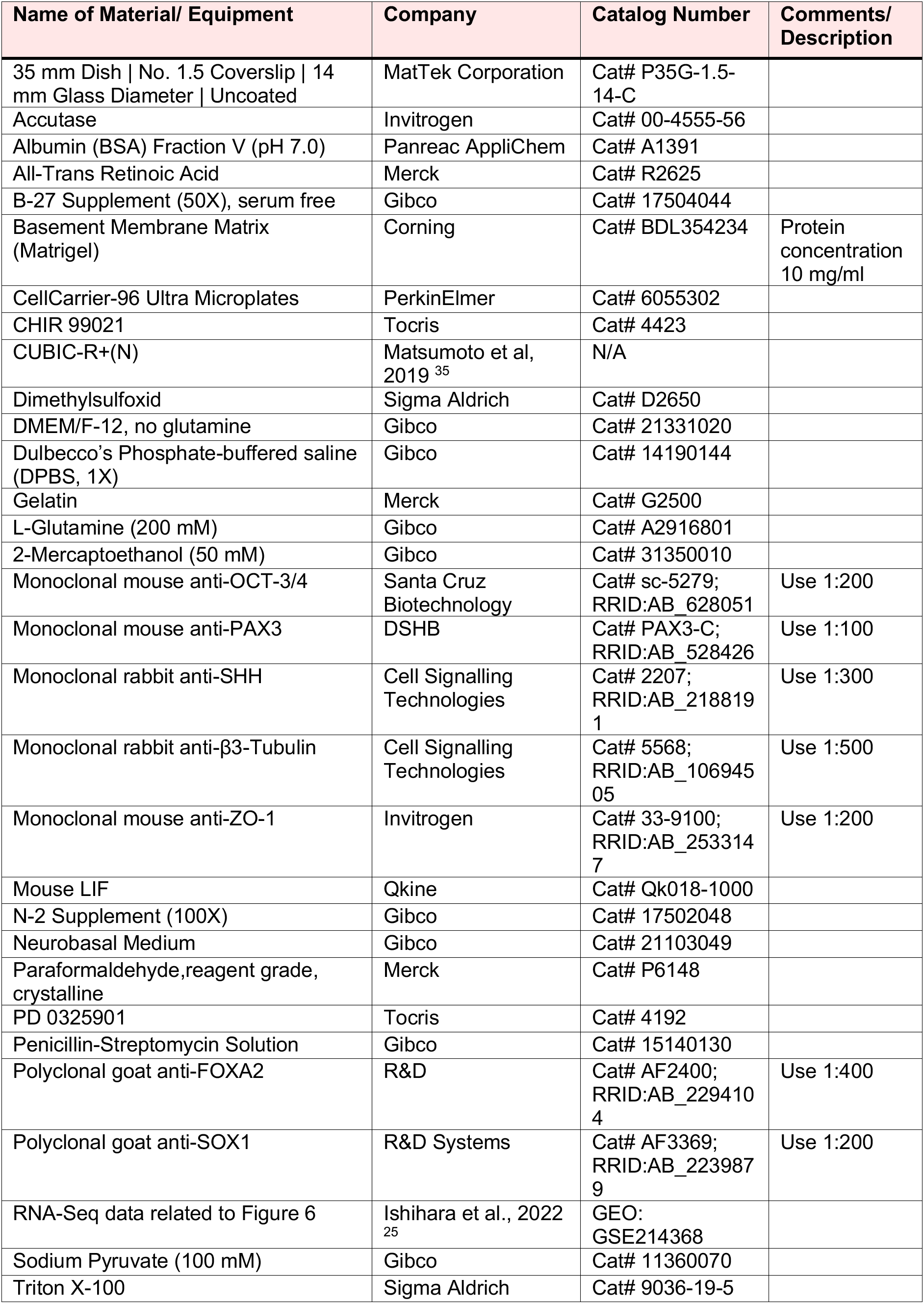

TABLE OF MATERIALS:
| Name of Material/ Equipment | Company | Catalog Number | Comments/ Description |
| --- | --- | --- | --- |
| 35 mm Dish \| No. 1.5 Coverslip \| 14 mm Glass Diameter \| Uncoated | MatTek Corporation | Cat# P35G-1.5-14-C |  |
| Accutase | Invitrogen | Cat# 00-4555-56 |  |
| Albumin (BSA) Fraction V (pH 7.0) | Panreac AppliChem | Cat# A1391 |  |
| All-Trans Retinoic Acid | Merck | Cat# R2625 |  |
| B-27 Supplement (50X), serum free | Gibco | Cat# 17504044 |  |
| Basement Membrane Matrix (Matrigel) | Corning | Cat# BDL354234 | Protein concentration 10 mg/ml |
| CellCarrier-96 Ultra Microplates | PerkinElmer | Cat# 6055302 |  |
| CHIR 99021 | Tocris | Cat# 4423 |  |
| CUBIC-R+(N) | Matsumoto et al, 2019<sup>35</sup> | N/A |  |
| Dimethylsulfoxid | Sigma Aldrich | Cat# D2650 |  |
| DMEM/F-12, no glutamine | Gibco | Cat# 21331020 |  |
| Dulbecco's Phosphate-buffered saline (DPBS, 1X) | Gibco | Cat# 14190144 |  |
| Gelatin | Merck | Cat# G2500 |  |
| L-Glutamine (200 mM) | Gibco | Cat# A2916801 |  |
| 2-Mercaptoethanol (50 mM) | Gibco | Cat# 31350010 |  |
| Monoclonal mouse anti-OCT-3/4 | Santa Cruz Biotechnology | Cat# sc-5279; RRID:AB_628051 | Use 1:200 |
| Monoclonal mouse anti-PAX3 | DSHB | Cat# PAX3-C; RRID:AB_528426 | Use 1:100 |
| Monoclonal rabbit anti-SHH | Cell Signalling Technologies | Cat# 2207; RRID:AB_2188191 | Use 1:300 |
| Monoclonal rabbit anti-β3-Tubulin | Cell Signalling Technologies | Cat# 5568; RRID:AB_10694505 | Use 1:500 |
| Monoclonal mouse anti-ZO-1 | Invitrogen | Cat# 33-9100; RRID:AB_2533147 | Use 1:200 |
| Mouse LIF | Qkine | Cat# Qk018-1000 |  |
| N-2 Supplement (100X) | Gibco | Cat# 17502048 |  |
| Neurobasal Medium | Gibco | Cat# 21103049 |  |
| Paraformaldehyde, reagent grade, crystalline | Merck | Cat# P6148 |  |
| PD 0325901 | Tocris | Cat# 4192 |  |
| Penicillin-Streptomycin Solution | Gibco | Cat# 15140130 |  |
| Polyclonal goat anti-FOXA2 | R&D | Cat# AF2400; RRID:AB_2294104 | Use 1:400 |
| Polyclonal goat anti-SOX1 | R&D Systems | Cat# AF3369; RRID:AB_2239879 | Use 1:200 |
| RNA-Seq data related to Figure 6 | Ishihara et al., 2022<sup>25</sup> | GEO: GSE214368 |  |
| Sodium Pyruvate (100 mM) | Gibco | Cat# 11360070 |  |
| Triton X-100 | Sigma Aldrich | Cat# 9036-19-5 |  |

## ACKNOWLEDGMENTS

We would like to thank present and past members of the Tanaka lab for their support and discussions in the development of this protocol. Especially, we thank A. Pelzl for critically reading the manuscript, giving input and helping with schematics, and to E. Chatzidaki for support. This publication was funded by the Austrian Science Fund (FWF): “Stand Alone publication project number PUD 25”. Additionally, this project has received funding from the European Research Council (ERC) under the European Union’s Horizon 2020 research and innovation programme (grant agreement ERC AdG 742046), and from the Austrian Science Fund (FWF) [10.55776/ F7803-B] (Stem Cell Modulation). For the purpose of Open Access, the authors have applied a CC BY public copyright license to any Author Accepted Manuscript (AAM) version arising from this submission.

